# CAGEfightR: Cap Analysis of Gene Expression (CAGE) in R/Bioconductor

**DOI:** 10.1101/310623

**Authors:** Malte Thodberg, Axel Thieffry, Kristoffer Vitting-Seerup, Robin Andersson, Albin Sandelin

## Abstract

We developed the CAGEfightR R/Biconductor-package for analyzing CAGE data. CAGEfightR allows for fast and memory efficient identification of transcription start sites (TSSs) and predicted enhancers. Downstream analysis, including annotation, quantification, visualization and TSS shape statistics are implemented in easy-to-use functions. The package is freely available at https://bioconductor.org/packages/CAGEfightR

## Introduction

Transcription Start Sites (TSSs) are central to gene regulation research. Cap Analysis of Gene Expression (CAGE) is a popular platform for genome-wide identification of TSSs, based on sequencing the first 20-30 bp of capped full-length RNAs, called CAGE tags^1^. Mapped to a reference genome, CAGE tags identify the location and measure the expression level of TSSs independent of reference transcript models^2^. They can also predict active enhancers based on bidirectional transcription initiation of enhancer RNAs (eRNAs)^3^.

Currently available tools for analyzing CAGE data (e.g. ^4–6^) were developed for older versions of the CAGE-protocol and/or are stand-alone tools. Here, we present the CAGEfightR R/Bioconductor package, which makes Bioconductor packages for analyzing RNA-Seq, ChIP-Seq and microarrays available for CAGE analysis. Compared to existing Bioconductor CAGE packages (e.g. CAGEr^7^, TSRchitect (http://doi.org/10.18129/B9.bioc.TSRchitect)), CAGEfightR uniquely offers enhancer prediction, novel methods for robust tag clustering, annotation and visualization, all implemented using standard Bioconductor classes.

## Methods

CAGEfightR functionality is illustrated using CAGE data from mouse lung tissue ^8^ and HeLa cells^9^; see Supplementary material.

## Results

The input for a CAGEfightR analysis is BigWig-files counting the occurrence of 5’ ends of CAGE tags at individual genomic bp, referred to as CAGE TSSs(CTSSs)^2^. CAGEfightR uses sparse matrices to efficiently store and manipulate large CTSS datasets using little memory; CAGEfightR can analyze tens of samples on a normal laptop, and hundreds of samples on a typical server. CAGEfightR analyses CAGE data at three levels: CTSS-, cluster- and gene-level (**Fig.1A**), described below. CAGEfightR provides efficient functions for importing multiple CTSS-files, quantifying and normalizing CTSSs counts to Tags-Per-Million(TPM), which can be summed across libraries to yield a pooled CTSS signal. Nearby CTSSs are commonly grouped into clusters for downstream analyses: CAGEfightR can find unidirectional tag clusters (TCs) for gene TSS identification and bidirectional clusters (BCs) for enhancer prediction.

**Fig. 1.**
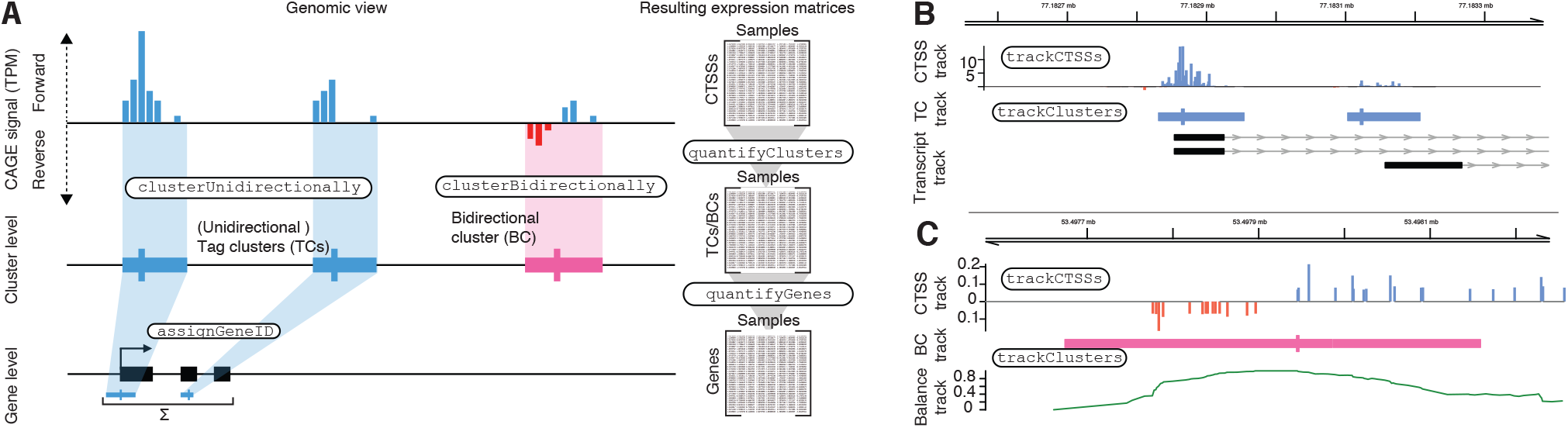
Conceptual overview. Boxed names indicate CAGEfightR functions. **A: CAGEfightR analyzes CAGE data at multiple levels.** Pooled CTSSs (top) can be used to find clusters (middle), which can be assigned to annotated genes (bottom). Each level is associated with an expression matrix (right). **B: Genome browser example of unidirectional tag clusters (TCs).** From the top, pooled CTSSs, TCs (peak indicated in thick) and transcript models. **C: Genome browser example of bidirectional cluster (BC)/predicted enhancer.** From the top, pooled CTSSs (as B), BCs (optimally balanced midpoint in thick) and calculated balance signal used for BC identification.

TCs are found by searching for CTSSs with a signal above a chosen threshold, and then merging nearby CTSS on the same strand into clusters (**Fig.1A-B**). The threshold can be tuned to avoid excessive merging of TSSs due to many singleton CTSSs and/or individual CTSSs can be discarded prior to clustering if they are not detected in a certain number of samples.

Unlike using an initial set of TCs as in ^3^, CAGEfightR scans the entire genome for BCs directly: upstream and downstream pooled CTSSs are quantified for every genomic position. Next, the Bhattacharyya Coefficient^10^ is used to quantify the departure of the observed CTSS signal from perfect bidirectionality (**Fig.S1A**). Sites with a balance score above a threshold are identified, and nearby sites are merged into BCs (**Fig.1C**), which can be used for enhancer prediction. CAGEfightR-predicted enhancers have similar enrichment for DNase hypersensitive sites and chromatin states in Hela cells as ^3^ (**Fig.S1B**).

A set of TCs and/or BCs (**Fig.1A**) can be quantified across all samples to yield a cluster-level expression matrix, which can be used as input to other Bioconductor expression analysis tools. It is often useful to annotate CAGE clusters in relation to known transcript models. Because of transcript complexity, clusters may simultaneously overlap annotated TSSs, exons and introns of different transcripts. CAGEfightR uses a hierarchical approach where conflicting annotations are resolved hierarchically, i.e. overlap to annotated TSS are prioritized vs. 5’ UTR regions, etc. (**Fig.S1C**). This annotation scheme allows for easy identification of known and novel TSSs (**Fig.S1C**).

CAGEfightR includes functions for calculating statistics for identifying broad and sharp TSSs distributions (the interquartile range and TC position entropy (**Fig.S1D**)), and a framework for implementing custom functions for user-supplied shape statistic.

In order to capitalize on existing data and tools contingent on gene-level expression, it is sometimes useful to measure gene expression using CAGE data. CAGEfightR includes functions for summarizing TC expression within genes to obtain a genelevel expression matrix (**Fig.1A**).

Another use of gene models is the study of alternative TSSs and differential TSS usage. CAGE can filter lowly transcribed TCs prior to analyses, by discarding TCs contributing with e.g. <10% of total gene expression.

## Conclusion

CAGEfightR is an R/Bioconductor package for the downstream analysis of CAGE data. CAGEfightR is the first single framework that robustly detects, quantifies, annotates and visualizes TSSs and enhancers from CAGE data in a manner that is highly compatible with other Bioconductor packages. The memory efficient and scalable implementation allows CAGEfightR to be used on datasets ranging from small-scale experiments to consortia-level projects. While developed for CAGE data, CAGEfightR can analyze any similar type of sparse genomic data, e.g. PEAT, RAMPAGE, CapSeq, etc., and with some modification also tag-based transcription assays, e.g. PRO-Seq, GRO-Seq and NET-Seq.

## Acknowledgements

We would like to acknowledge past and present members of the Sandelin Lab for help and input on CAGE data analysis.

## Supplementary material for CAGEfightR: Cap Analysis of Gene Expression (CAGE) in R/Bioconductor by Thodberg *et al*

### Supplementary Methods

Datasets were obtained as described in Methods. For mouse samples, CAGEfightR was run on default settings. Transcript models and genomic sequence were obtained from Bioconductor GenomicFeatures^1^ annotation packages *TxDb.Mmusculus.UCSC.mm9.knownGene* and *BSgenome.Hsapiens.UCSC.hg19*.

For immortalized cell lines, Andersson *et al* enhancers were obtained by running scripts from https://github.com/anderssonrobin/enhancers, using CAGEfightR TCs as input. All analyzes were run with default parameters. Only enhancers located outside of known transcripts (as in **Fig.S1C**) were kept for analysis based on annotation packages *TxDb.Hsapiens.UCSC.hg19.knownGene*. HeLa DNase peaks^2^ and chromatin states^3^ were obtained via Bioconductor *AnnotationHub* records: AH30755 & AH46970.

---

**Supplementary Figure 1:**
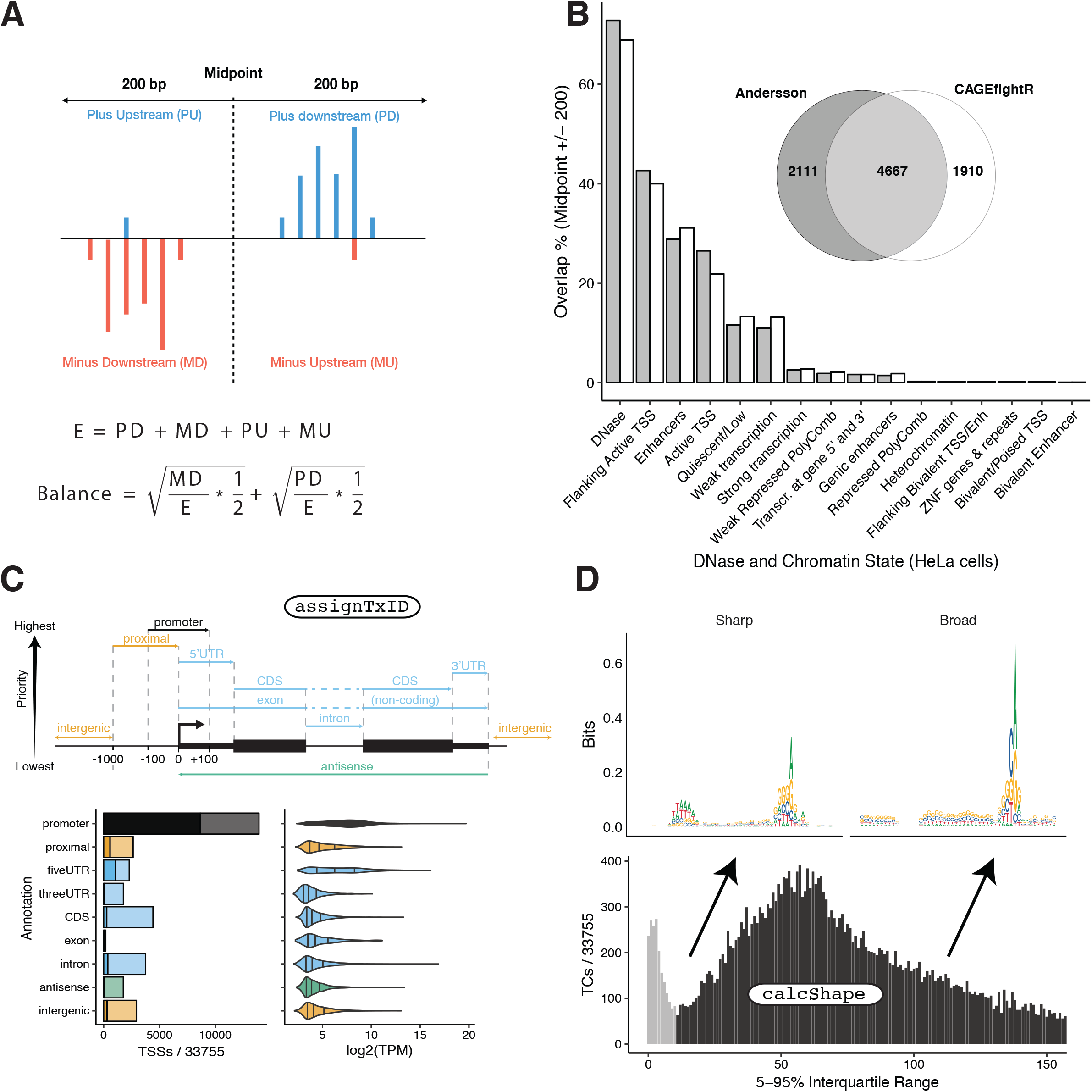
**A: Using the Bhattacharyya Coefficient to find bidirectional clusters.** CAGEfightR scans every position of the genome using the Bhattacharyya Coefficient on pooled CTSSs in +/-200 bp windows, resulting in a genome wide balance score (see **Fig.1C** bottom track) **B: Comparison between CAGEfightR-defined enhancers and enhancers defined by Andersson et al:** Top-right: Venn diagram showing numbers and overlap of enhancers detected by Andersson et al (grey) and CAGEfightR (white). Bottom: X-axis shows HeLa cell chromatin states and DNase peaks from the ENCODE project. Y-axis shows the overlap of Andersson et al enhancers (grey) and CAGEfightR (white) as percentage of total the number of called enhancers. Overlaps are calculated using enhancer balance midpoints +/-200 bp. **C: Hierarchical annotation of clusters.** Top: Cartoon of hierarchical annotation sheme in relation to known transcripts (bp distances can be modified by the user). Annotations towards the top have higher priority over ones toward the bottom. Bottom: Example analysis based on assignTxID, plotting the counts and expression of annotated TCs. Y-axis shows the annotation categories. Left panel shows the number of TCs in each category expressed at more than 1 TPM (light) or 10 TPM (dark) in more than 2 samples. Right panel: X-axis shows the distribution of expression of each category **A: Example of shape statistic calculation enabled by CAGEfightR.** Bottom: Histogram showing 5-95% interquartile range of tag clusters (TCs), with sharp (<10 bp) and broad (>10 bp) marked. X-axis is cut at 150 bp.Top: As a check whether interquartile range calculation recapitulates known features of broad and sharp TCs, sequence logos of DNA sequences underlying TCs were aligned based on their TC peaks. As expected, sharp TCs feature TATA boxes while broad clusters have a more prominent INR.

## References

1. Takahashi, H., Lassmann, T., Murata, M. & Carninci, P. 5’ End-Centered Expression Profiling Using Cap-Analysis Gene Expression and Next-Generation Sequencing. Nat. Protoc. 7, 542–61 (2012).

2. Carninci, P. et al. Genome-wide analysis of mammalian promoter architecture and evolution. Nat. Genet. 38, 626–35 (2006).

3. Andersson, R. et al. An atlas of active enhancers across human cell types and tissues. Nature 507, 455–61 (2014).

4. Ohmiya, H. et al. RECLU: A pipeline to discover reproducible transcriptional start sites and their alternative regulation using capped analysis of gene expression (CAGE). BMC Genomics 15, 1–15 (2014).

5. Hasegawa, A., Daub, C., Carninci, P., Hayashizaki, Y. & Lassmann, T. MOIRAI: A compact workflow system for CAGE analysis. BMC Bioinformatics 15, 1–7 (2014).

6. Frith, M. C. et al. A code for transcription initiation in mammalian genomes. Genome Res. 18, 1–12 (2008).

7. Haberle, V., Forrest, a. R. R., Hayashizaki, Y., Carninci, P. & Lenhard, B. CAGEr: precise TSS data retrieval and high-resolution promoterome mining for integrative analyses. Nucleic Acids Res. 1–11 (2015). doi:10.1093/nar/gkv054

8. Bornholdt, J. et al. Identification of Gene Transcription Start Sites and Enhancers Responding to Pulmonary Carbon Nanotube Exposure in Vivo. ACS Nano 11, 3597–3613 (2017).

9. Andersson, R. et al. Nuclear stability and transcriptional directionality separate functionally distinct RNA species. Nat. Commun. 5, 1–10 (2014).

10. Bhattacharyya, A. On a measure of divergence between two statistical populations defined by their probability distribution. Bull. Calcutta Math. Soc. 99–100 (1943).

## Supplementary References

1. Lawrence, M. et al. Software for Computing and Annotating Genomic Ranges. PLoS Comput. Biol. 9, 1–10 (2013).

2. Thurman, R. E. et al. The accessible chromatin landscape of the human genome. Nature 489, 75–82 (2012).

3. Dunham, I. et al. An integrated encyclopedia of DNA elements in the human genome. Nature 489, 57–74 (2012).

